# Feminisation of complex traits in *Drosophila melanogaster* via female-limited X chromosome evolution

**DOI:** 10.1101/2020.04.01.019737

**Authors:** Katrine K. Lund-Hansen, Jessica K. Abbott, Edward H. Morrow

## Abstract

A handful of studies have investigated sexually antagonistic constraints on obtaining sex-specific fitness optima, though exclusively through male-genome-limited evolution experiments. In this paper, we established a female-limited X chromosome evolution experiment, where we used an X chromosome balancer to enforce the inheritance of the X chromosome through the matriline, thus removing exposure to male selective constraints. This approach eliminates the effects of sexually antagonistic selection on the X chromosome, permitting evolution towards a single sex-specific optimum. After multiple generations of selection, we found strong evidence that body size and development time had moved towards a female-specific optimum, whereas reproductive fitness and locomotion activity remained unchanged. The changes in body size and development time are consistent with previous results, and suggest that the X chromosome is enriched for sexually antagonistic genetic variation controlling these traits. The lack of change in reproductive fitness and locomotion activity could be due to a number of mutually non-exclusive explanations, including a lack of sexually antagonistic variance on the X chromosome or confounding effects of the use of the balancer chromosome. This study is the first to employ female-genome-limited selection and adds to the understanding of the complexity of sexually antagonistic genetic variation.

## Introduction

Genetic conflict can occur in dioecious species when the shared genome cannot fully accommodate the evolutionary interests of both females and males. There is, therefore, a theoretical expectation that such genetic constraints will limit each sex from reaching its respective phenotypic optimum. Furthermore, alleles that increase fitness in one sex can be directly detrimental to the other, i.e. sexually antagonistic alleles (Parker 1979; Rice and Chippindale 2002; Bonduriansky and Chenoweth 2009). Sexually antagonistic selection, the evolutionary process where selection acts in opposing directions on the two sexes, has been likened to a genomic tug-of-war between the sexes (Rice and Chippindale 2001). Sexually antagonistic selection has been shown to be common in wild populations across many different vertebrate and invertebrate taxa (Cox and Calsbeek 2009) and there is evidence that sexually antagonistic genetic variation is substantial within *Drosophila melanogaster* (Chippindale et al. 2001; Gibson et al. 2002; Innocenti and Morrow 2010; Ruzicka et al. 2019).

A valuable way to test the presence of sexually antagonistic selection and its impact on shared phenotypic traits is sex-limited experimental evolution, where natural selection is limited, by the investigator, to one sex (Rice 1996). Previous evolution experiments have shown that it is possible to alter selection on the genome to favour one sex over another, thereby allowing trait values to evolve towards the optimum of the selected sex. These selection experiments were achieved either through genome-limited selection (Rice 1996, 1998; Prasad et al. 2007; Abbott et al. 2013) or the removal or limitation of sexual selection (Holland and Rice 1999; Pitnick et al. 2001; Morrow et al. 2008; Nandy et al. 2013; Innocenti et al. 2014). So far, genome-limited evolution experiments have been restricted to allow selection in males only. It is therefore a natural extension to undertake a female-limited evolution experiment to test sexual conflict theory using a complementary design.

In this paper, we test the hypothesis that eliminating male selection on the X chromosome via a female-limited X chromosome (FLX) evolution experiment would release sexually antagonistic standing genetic variation from the intersexual tug-of-war. By using an X chromosome balancer, which does not recombine with its homolog, we enforced female-limited transmission of the X chromosome. We chose to limit selection to the X chromosome only, as both theory (Rice 1984) and some empirical evidence (Gibson et al. 2002; Innocenti and Morrow 2010) indicate that the X chromosome is enriched for sexually antagonistic variance for fitness. Still, the evidence for the X chromosome being a “hot spot” for sexually antagonistic variance is not unequivocal (Fry 2010; Ruzicka et al. 2019).

We predicted that expressing the experimentally feminised X chromosomes would shift phenotypic trait values towards female-specific optima in both sexes. We chose to investigate phenotypic traits that have previously shown to change under male-limited (ML) evolution experiments, such as reproductive success/fitness, development time and body size (Prasad et al. 2007). We predicted an increase in female reproductive fitness at the cost of male reproductive fitness, an increase in body size and a decrease in development time. We also tested locomotion activity, as it has been shown to be a sexually antagonistic trait (Long and Rice 2007). We expected to find a decreased female locomotory activity in the FLX selected females. Some, but not all, of these predictions were supported by our results.

## Methods

### Fly stocks

We used the LH_M_ stock as the base population for the evolution experiment. LH_M_ comes from a large laboratory-adapted population collected in central California in 1991 and founded from 400 inseminated females (Rice et al. 2005). This stock has been maintained in our lab for over 500 generations under the standard LH_M_ culturing protocol: at 25°C, 12-12 light-dark cycle, 60% relative humidity, and fed on cornmeal-molasses-yeast medium (Rice et al. 2005).

For the experimental evolution we used an X chromosome balancer stock, FM7a (In(1)B^1^/sc^8^/v^Of^/w^a^/y^31d^), obtained from Adam Chippindale, Queen’s University. We backcrossed the FM7a stock into LH_M_ for 12 generations, by crossing FM females to LH_M_ males. To ensure that the mitochondrial genetic background was also derived from LH_M_ we crossed LH_M_ females to FM males for one generation. Each cross was conducted with 672 breeding adults (♀16:16♂ per vial) in 42 vials.

### Experimental setup

After 12 generations of backcrossing, we established three treatments (see below) in four replicate populations, giving a total of 12 populations. Each treatment was maintained as an adult breeding population of 448 individuals (♀16:16♂ per vial) in 14 vials with a minimum of 2,100 offspring.

#### Female-limited X chromosome (FLX) treatment

To ensure female-limited evolution of the X chromosome, we crossed heterozygote (FM/X) females to FM/Y males. Thus, females always inherit the evolving X chromosome from their mother and an FM balancer from their father (Figure 1). FM/X females were collected as virgins every generation to control the female-limited inheritance pattern of the X chromosome.

**Figure 1:**
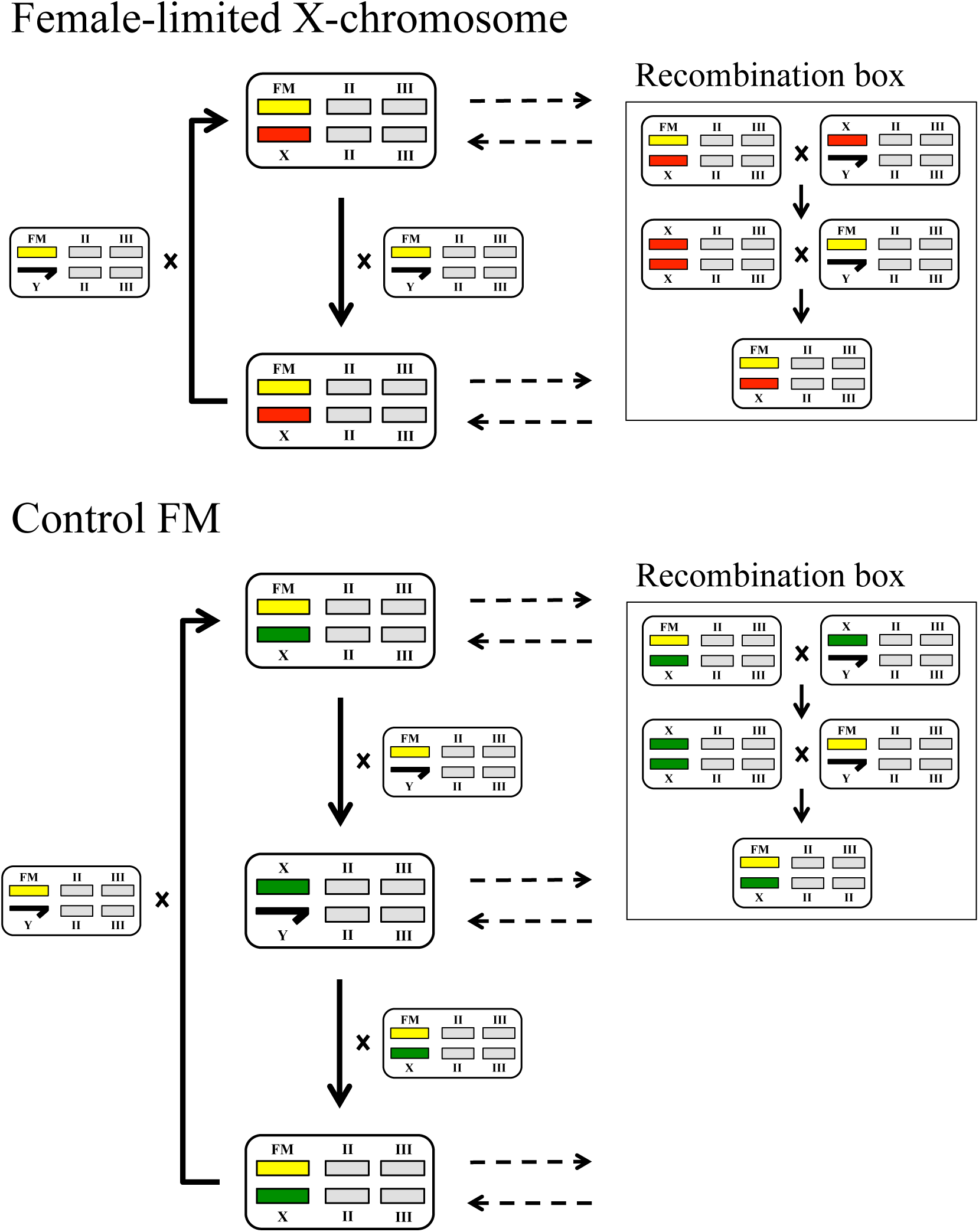
Protocol for the Female-limited X chromosome (FLX) evolution experiment, the FM balancer control (CFM), and the recombination box for both. All chromosomes are depicted by different coloured rectangles: light grey (autosome II and III), yellow (FM balancer), red (FLX X chromosome), and green (CFM X chromosome). The Y chromosome is depicted as the black half-arrow. For additional details see *Evolution experiment protocol* in Methods.

#### Control FM (CFM) treatment

We included a methodological control group to control for unforeseen effects of adaptation to the presence of the FM balancer in the FLX populations. We wanted the X chromosome to be present with the FM balancer but to follow the average inheritance pattern of X chromosomes in wild type populations by being present in males one-third of the time. Thus, every third generation FM/X females were crossed to CFM X/Y males, so the X chromosome was passed from father to daughter (Figure 1). As with the FLX treatment, all females were collected as virgins.

#### Control wild type (Cwt) treatment

A group of wild-type flies was maintained under the same experimental conditions as the FLX treatment (virgin collection, smaller population size), but without sex-limited selection or the FM balancer. We were thereby able to control for the experimental protocol itself and for any effects that may be caused by a reduction in effective population size.

With this set-up, we can expect that traits where the FLX and CFM treatments both differ from the Cwt treatment, but are similar to each other, are likely a result of adaptation to the FM balancer. In contrast, traits where the FLX treatment differs from the two other treatments, but the CFM and Cwt treatments are similar to each other, are likely a result of female-limited selection on the X chromosome.

#### Recombination box

Because the X chromosome does not recombine with the FM balancer, the small effective population size of the X chromosome could lead to an accumulation of deleterious mutations due to genetic drift and/or hitchhiking (Futuyma 2009). This finite population size could also slow selection on the experimental X chromosome due to the Hill-Robertson effect, where selection on one locus can affect the efficiency of selection on a second locus, especially if they are in linkage, thereby reducing the overall of efficiency of selection (Hill and Robertson 1966). Rice (1996) showed that allowing recombination in a small percentage of the experimental population could reduce these problems. To allow this, every generation 16 FM/X flies (4%) were removed from each of the evolving FLX populations to go through recombination. These 16 FM/X flies were crossed to X/Y males from the same populations, so the two X chromosomes could recombine in X/X females in the next generation. The X/X females were then crossed to FM/Y males, so the recombined X chromosomes could be returned the evolving populations, as FM/X females. Using this method, one recombined X chromosome was added to each vial every generation, so 14 FM/X flies (3%) were returned to the evolving populations (see the recombination box for FLX in Figure 1). As the X chromosome in CFM is only co-located with the FM balancer in females, each generation the same number of flies were removed to undergo recombination as in the FLX treatment (see the recombination box for CFM in Figure 1).

### Expression of the X chromosome in females and males

For all phenotypic assays, we wanted to express the evolved X chromosomes in both sexes without the FM balancer, as it was only a methodological device. Because females carry two X chromosomes, we decided to include two female types: females with two evolving X chromosomes (2X), and females with one evolving and one wild type X chromosome (1X). It was not clear *a priori* which female type was the most appropriate to investigate because although the effects of the selection treatment could potentially be more apparent in 2X females, there was also the risk that they could be inbred on the X. We were also interested in being able to test hypotheses about dominance variance on the X. One generation before each assay we crossed FLX FM/X females to either FLX X/Y males (to produce FLX-2X females and FLX males) or Cwt males (to produce FLX-1X females). As the CFM treatment females also always carried an FM balancer, for the phenotypic assays we also include two female types as well as CFM X/Y males. One generation before each assay we crossed CFM FM/X females to either CFM X/Y males (to produce CFM-2X females and CFM males) or Cwt males (to produce CFM-1X females).

### Development time

We collected females as virgins on day 10 after oviposition and crossed them to males on day 12. For each type and replicate population, we sat up five vials with 16 pairs in each. After the flies had interacted for two days, we moved the females to fresh vials where they oviposit for 18 hours. We reduced the egg number to 150-200 eggs after discarding the adult flies. On day nine after oviposition, we started vial observation. The vials were subsequently observed approximately once every six hours until no flies had eclosed for seven consecutive observations. At every observation we cleared the vials of eclosed flies and the flies were scored for sex and phenotype. This assay was performed at generation 43.

### Female fecundity assay

We measured female fecundity as the number of eggs laid by a female during an 18 hour period, which matches the period normally available to females from the base population (Rice et al. 2005). We collected five adult virgin females (day 10 after oviposition) and combined them with ten competitor virgin females and 15 males in a yeasted vial on day 12. The competitor females and males were from the outbred LH_M_ population homozygous for the visible brown eye (*bw*) genetic marker (LH_M_-*bw*) that is recessive to the wild-type red eye-colour allele. After two days, we isolated the five target females in individual test tubes with food medium for 18 hours to oviposit. The females were then discarded, and we froze the test tubes so we could count the eggs without larvae eclosing. This assay was done at two different time points, generation 15 and 41. At generation 15, we carried out the assay in 10 replicates per female type per replicated population, thereby providing fitness estimates based on data from 200 individuals per type. At generation 41, we did the assay in 12 replicates per female type per replicated population, thereby providing fitness estimates based on data from 240 individuals per type. We also used this protocol to measure the fitness of females with the FM balancer. The FM balancer fitness assay was done in two blocks, with five replicates of six female types in each block, thereby providing fitness estimates based on data from 200 individuals per type. We calculated relative fecundity by dividing the fitness for each replicate by the maximum fitness across all replicates.

### Locomotion activity

We adapted the protocol from Long and Rice (2007) to measure locomotion activity. On day 12 after oviposition, we collect five adult non-virgin flies of the same sex and type. The flies were left for 24 hours to ensure that they had fully recovered from CO_2_ anaesthesia during collection. We drew a large rectangle partitioned into eighths on each vial and randomly chose one fly within one section for observation. The focal fly was observed for 3s and we noted if the fly was walking around (active) or not. Each vial was observed over 10 separate sessions on the same day. We carried out this assay in five replicates per female and male type per replicate population (for a total of 20 vials) at generation 123.

### Male fitness assay

We measured male fitness as the proportion of live offspring sired by males carrying the target X chromosome during competition using a standard LH_M_ eye-marker protocol (Rice et al. 2005; Abbott et al. 2013). We combined five adult target males (day 13 after oviposition) with ten competitor LH_M_-*bw* males and 15 virgin LH_M_-*bw* females in a yeasted vial. After two days, we isolated the 14 females in individual tubes with food medium for 18 hours to oviposit. The females were then discarded, and the test tubes were left under standard LH_M_ conditions for 12 days. We counted the adult offspring from each tube and scored them for eye-colour to assign paternity between target and competitor males. Since the wild-type (red) eye-colour allele is dominant to the *bw* allele, offspring with a red eye-colour can be assigned to the target males. As with the female assay, we carried out this assay at two different time points. At generation 18, we carried out the assay in 10 replicates for each of the three male types, thereby providing fitness estimates based on data from 200 individuals per type. At generation 39 and 40, we carried out the assay in two blocks, with six replicates of the three male types in each block, thereby providing fitness estimates based on data from 240 individuals per type. We also used this protocol to measure the fitness of males with the FM balancer. We performed the FM balancer fitness assay in two blocks, with five replicates of six male types in each block, thereby providing fitness estimates based on data from 200 individuals per type. We calculated relative proportion of paternity share by dividing the fitness for each replicate by the maximum fitness across all replicates.

### Thorax size

We estimated body size using measurements of thorax length. On day 13 from oviposition, we collected 20 flies from each type and placed them in 95% ethanol. We measured air-dried flies using a Nikon SMZ800 dissecting microscope at 63x magnification fitted with an eyepiece graticule. The flies were measured lying on their right side, from the start of the prescutum to the end of the scutellum. This assay was carried out at generations 18 and 72. To ensure repeatability of the measurements, one person measured all the flies and we presumed that any bias would be equalised over the four replicate populations.

### Statistical procedures

All statistical analyses were conducted in R version 3.4.4 (R Core Team 2018). Because we had two female types and only one male type within the FLX and CFM treatments (see Expression of the X chromosome in females and males), we decided to test if there was a significant phenotypic difference between the two types of females before comparing the two sexes. For every phenotypic assay we did a Student’s t-Test (*t*.*test*) between the two female types within each treatment (Table 1). As there was no significant effect for any comparison, we grouped the two female types together within treatment in the statistical analysis for comparisons of the two sexes (a separate analysis of dominance effects within females is currently in preparation). For all phenotypic assays we used population means as the unit of the analyses, as was done previously for the ML evolution experiment (Prasad et al. 2007). Results were qualitatively similar when analysed with population as a random effect rather than using population means (see supplementary information). We carried out fully factorial linear models for all analyses with treatment and sex as fixed factors. When applicable we included block as a random factor. For the FM fitness assay we analysed the two sexes separately because the FM genotypes are not directly comparable. For significance tests we performed an ANOVA (*anova)* analysis. For the locomotion activity assay, we tested the number of active and inactive flies using Generalized Linear Models (*glm*), with family set as binomial and treatment, sex, and their interaction as fixed effects. For significance tests we performed an ANOVA (*anova*) analysis. We made post-hoc pairwise comparisons using Least-squares means (*emmeans*) (Lenth 2016).

**Table 1:** Summary of the t-tests between the two FLX female types and between the two control FM (CFM) female types

| Source | *df* | *t* | *P* |
| --- | --- | --- | --- |
| Fitness, generation 15/18 |  |  |  |
| FLX 1X vs. 2X | 3.11 | 2.21 | 0.11 |
| CFM 1X vs. 2X | 4.58 | -0.69 | 0.53 |
| Fitness, generation 39/41 |  |  |  |
| FLX 1X vs. 2X | 5.56 | 0.91 | 0.40 |
| CFM 1X vs. 2X | 4.18 | 1.52 | 0.20 |
| Thorax size, generation 15/18 |  |  |  |
| FLX 1X vs. 2X | 4.19 | 1.17 | 0.30 |
| CFM 1X vs. 2X | 5.40 | -0.27 | 0.80 |
| Thorax size, generation 72 |  |  |  |
| FLX 1X vs. 2X | 4.00 | -1.78 | 0.15 |
| CFM 1X vs. 2X | 3.48 | 0 | 1 |
| Development time, generation 43 |  |  |  |
| FLX 1X vs. 2X | 5.25 | 0.28 | 0.79 |
| CFM 1X vs. 2X | 5.18 | -0.04 | 0.97 |
| Locomotion, generation 123 |  |  |  |
| FLX 1X vs. 2X | 13.99 | 0 | 1 |
| CFM 1X vs. 2X | 13.32 | 0 | 1 |

## Results

There was no significant effect of the interaction between treatment and sex on reproductive fitness at either generation 15/18 (Figure 2.A and Table 2) or generation 39/41 (Figure 2.B and Table 2). At generation 15/18 there were indications that carrying the FLX-selected X chromosome could increase female fitness at the detriment of male fitness as predicted. But after a further 20 generations these indications did not persist and there were no differences in fitness between the treatments.

**Table 2:** Summary of the results from ANOVA analysis of linear models.

| Source | *df* | F | $\chi^2$ | *P* |
| --- | --- | --- | --- | --- |
| Fitness, generation 15/18 |  |  |  |  |
| Treatments | 2 | 0.75 |  | 0.48 |
| Sex | 1 | $7.15e^{-06}$ | | 1.00 |
| Treatments × Sex | 2 | 2.38 | | 0.11 |
| Fitness, generation 39/41 |  |  |  |  |
| Treatments | 2 | 2.01 |  | 0.15 |
| Sex | 1 | 0.08 |  | 0.83 |
| Treatments × Sex | 2 | 0.39 | | 0.68 |
| Thorax size, generation 15/18 |  |  |  |  |
| Treatments | 2 | 128.83 | | $3.22e^{-14}$ |
| Sex | 1 | 2285.00 | | $< 2.2e^{-16}$ |
| Treatments × Sex | 2 | 2.39 | | 0.11 |
| Thorax size, generation 72 |  |  |  |  |
| Treatments | 2 | 20.95 | | $3.80e^{-06}$ |
| Sex | 1 | 519.11 | | $< 2.2e^{-16}$ |
| Treatments × Sex | 2 | 0.43 | | 0.66 |
| Development time, generation 43 |  |  |  |  |
| Treatments | 2 | 9.84 | | $6.58e^{-04}$ |
| Sex | 1 | 1.22 |  | 0.28 |
| Treatments × Sex | 2 | $3.20e^{-03}$ | | 1.00 |
| Locomotion, generation 123 |  |  |  |  |
| Treatments | 2 |  | 2.06 | 0.36 |
| Sex | 1 | | 14.95 | $1.10e^{-04}$ |
| Treatments × Sex | 2 | | 4.79 | 0.09 |
| FM female fitness assay, generation 127 |  |  |  |  |
| Treatments | 5 | 0.53 |  | 0.75 |
| FM male fitness assay, generation 127 |  |  |  |  |
| Treatments | 5 | 54.04 | | $< 2.2e^{-16}$ |

**Figure 2:**
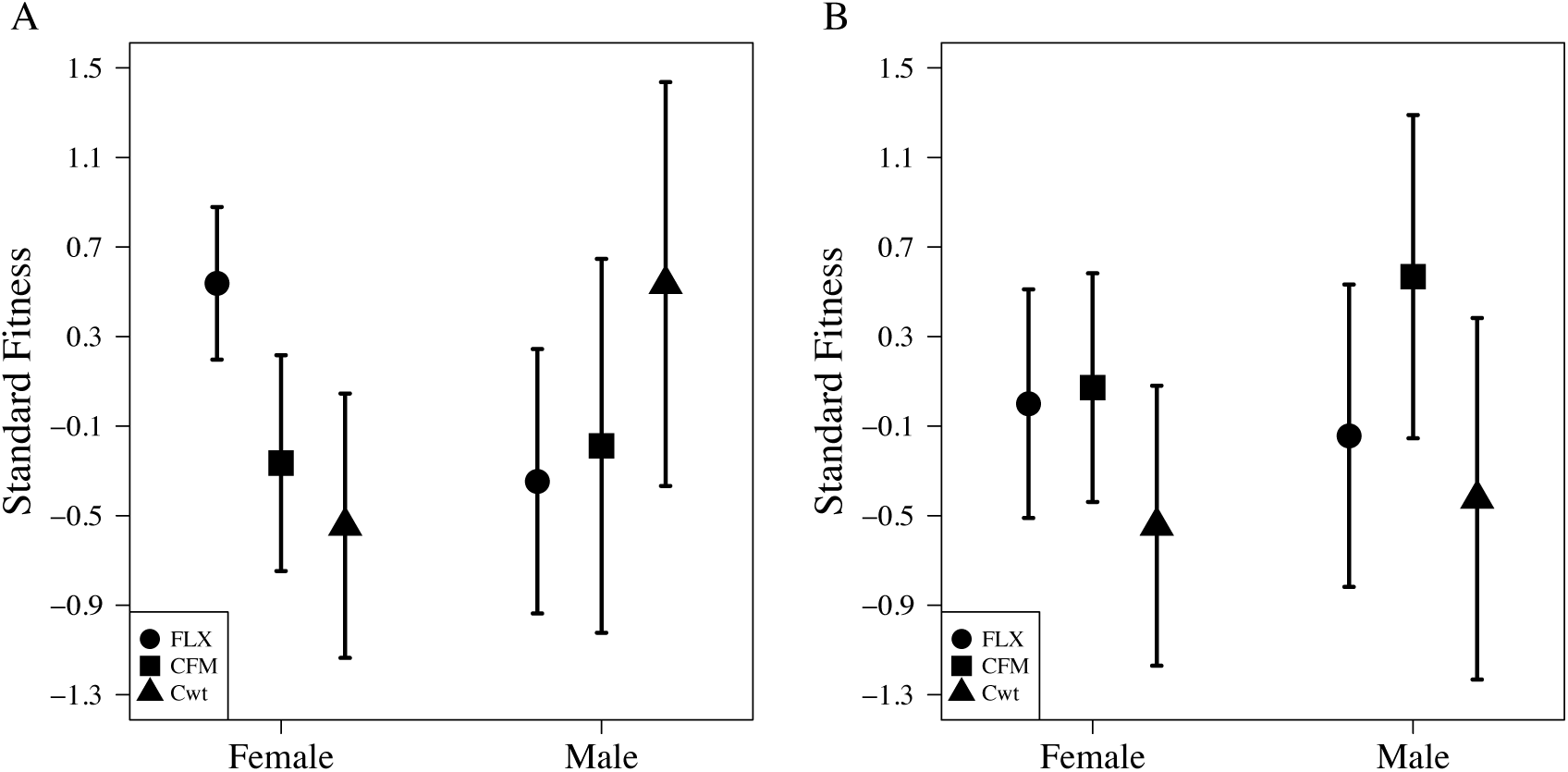
Standardised fitness at: (A) generation 15/18 and (B) generation 39/41. The selected or controlled X chromosomes are expressed in either females or males in all treatments. FLX: circle, control FM (CFM): square, and control wild-type (Cwt): triangle. Mean (±SE) average across the four replicate populations.

There was no significant effect of the interaction between treatment and sex on thorax size at generation 15/18 or at generation 72 (Figure 3 and Table 2). However there were significant main effects of sex and treatment at both time points (Table 2). At generation 15/18 FLX flies were significantly larger than the other two treatments in both sexes (Figure 3.A), however at generation 72, though the FLX were still nominally larger than the Cwt flies the difference was no longer significant (Figure 3.B). These results are consistent with the prediction that FLX flies would become larger during the evolution experiment.

**Figure 3:**
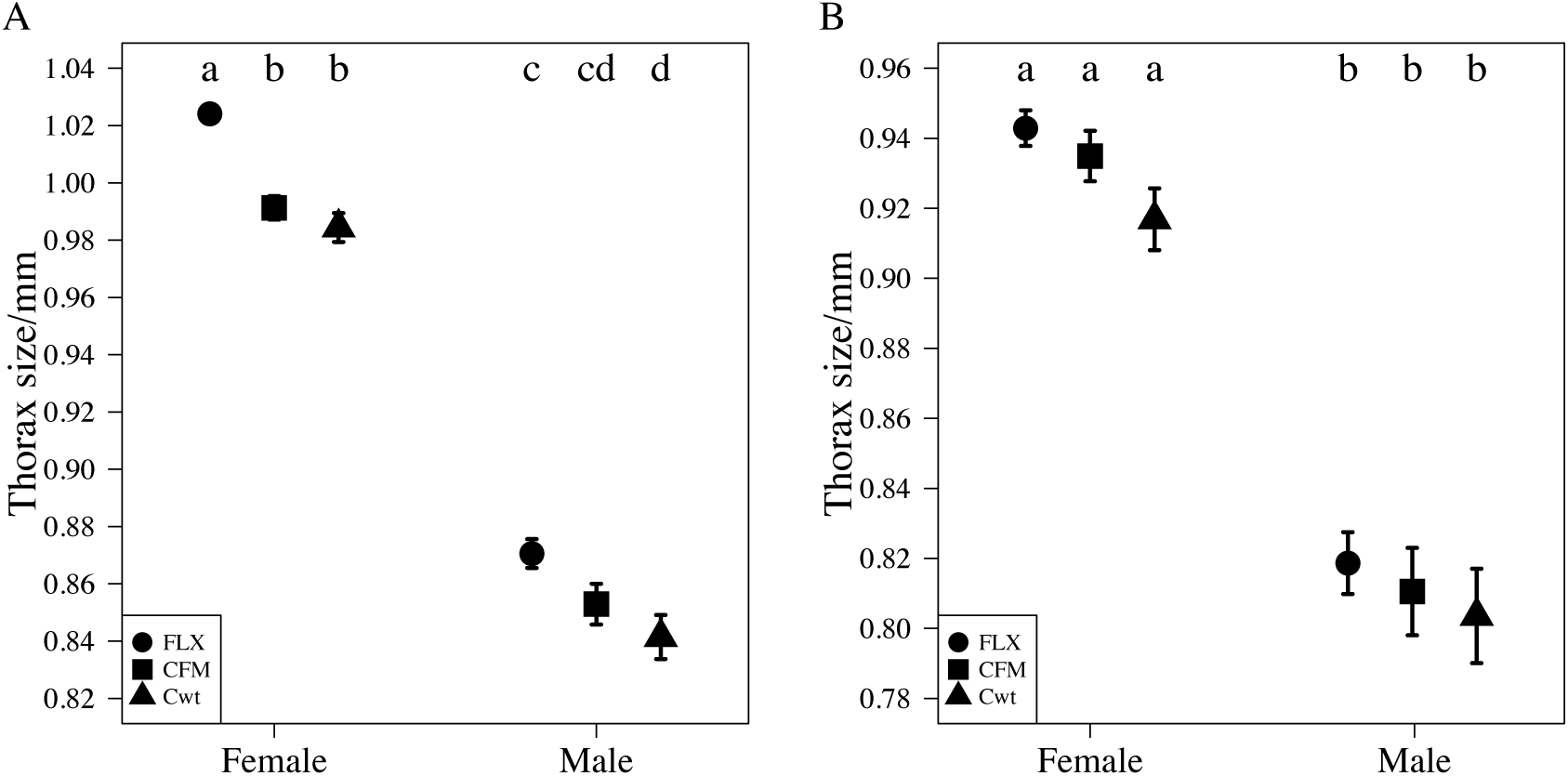
Thorax size in mm at: (A) generation 15 & 18 and (B) generation 72. The selected or controlled X chromosomes are expressed in either females or males in all treatments. FLX: circle, control FM (CFM): square, and control wild-type (Cwt): triangle. Mean (±SE) average across the four replicate populations. Letters indicates significance (Tukey HSD; *P* < 0.05).

There was no significant effect of the interaction between treatment and sex on development time, but there was a significant effect of treatment (Figure 4.A and Table 2). Both FLX females and males develop significantly faster than Cwt females and males, respectively (Figure 4.A). These findings are consistent with previous ML results, which showed that ML flies developed slower than controls (Prasad et al. 2007).

**Figure 4:**
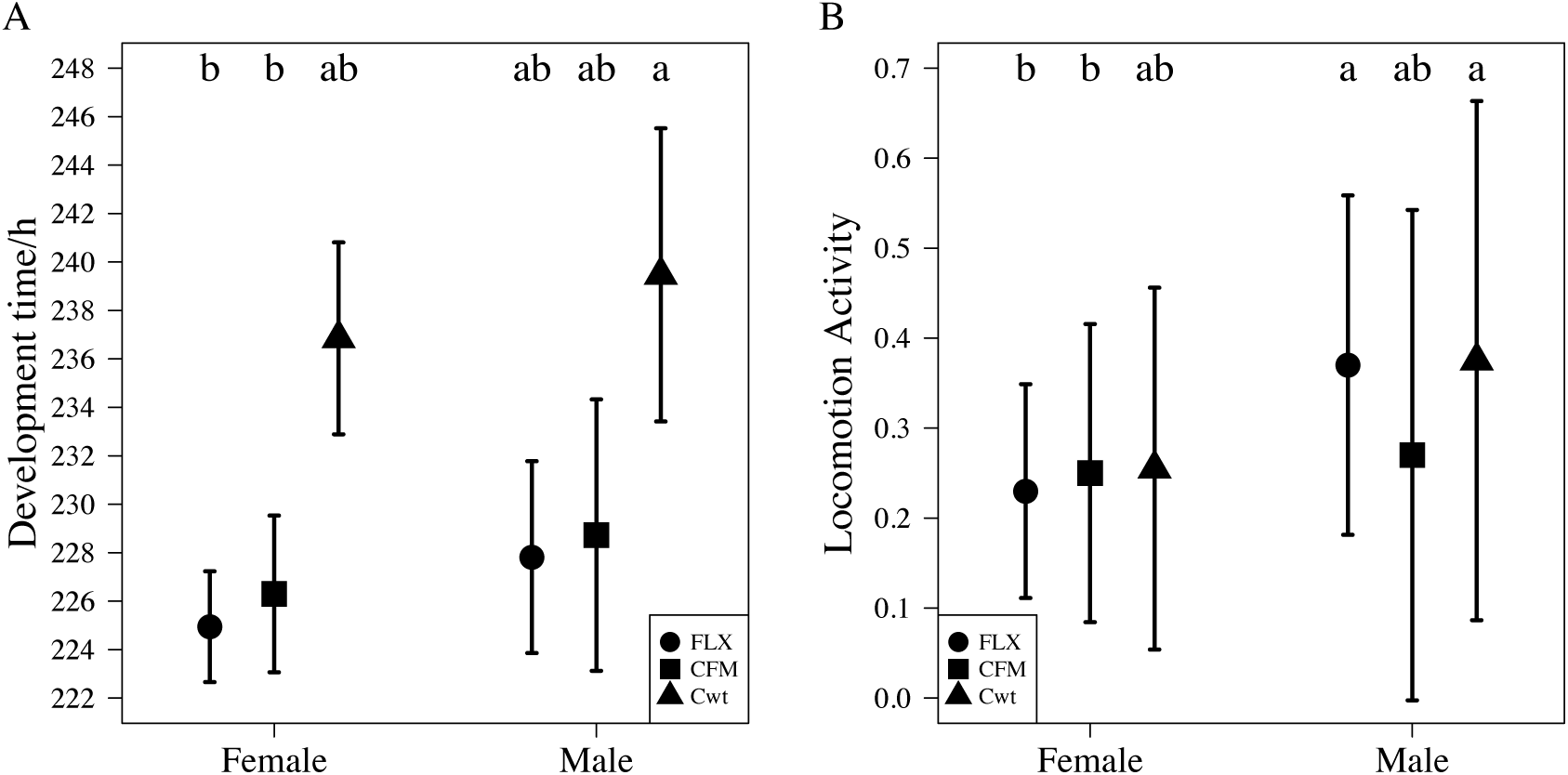
Behavioural assays of: (A) development time in hours at generation 43 and (B) locomotion activity at generation 123. The selected or controlled X chromosomes are expressed in either females or males in all treatments. FLX: circle, control FM (CFM): square, and control wild-type (Cwt): triangle. Mean (±SE) average across the four replicate populations. Letters indicates significance (Tukey HSD; *P* < 0.05).

There was a marginally significant effect of the interaction between treatments and sex and a significant effect of sex on locomotion activity (Figure 4.B and Table 2). However, FLX flies were not significantly less active than Cwt flies (Figure 4.B), which is not consistent with previous results showing that locomotory activity is a sexually antagonistic trait (Long and Rice 2007; Abbott et al. 2019).

It became clear from the phenotypic assays that the FM balancer did have unforeseen effects on fitness and locomotion, especially evident in males. We therefore decided to investigate what effect carrying the FM balancer actually had on the flies’ fitness. There was no significant difference in reproductive fitness between X/X and FM/X females (Figure 5.A and Table 2). There was a highly significant difference in reproductive fitness between X/Y and FM/Y males (Table 2). FM/Y males had significantly lower reproductive fitness than their X/Y counterparts (Figure 5.B) with a reduction in fitness of 41% for FLX FM/Y males, of 56% for CFM FM/Y males, and of 58% for Cwt FM/Y males.

**Figure 5:**
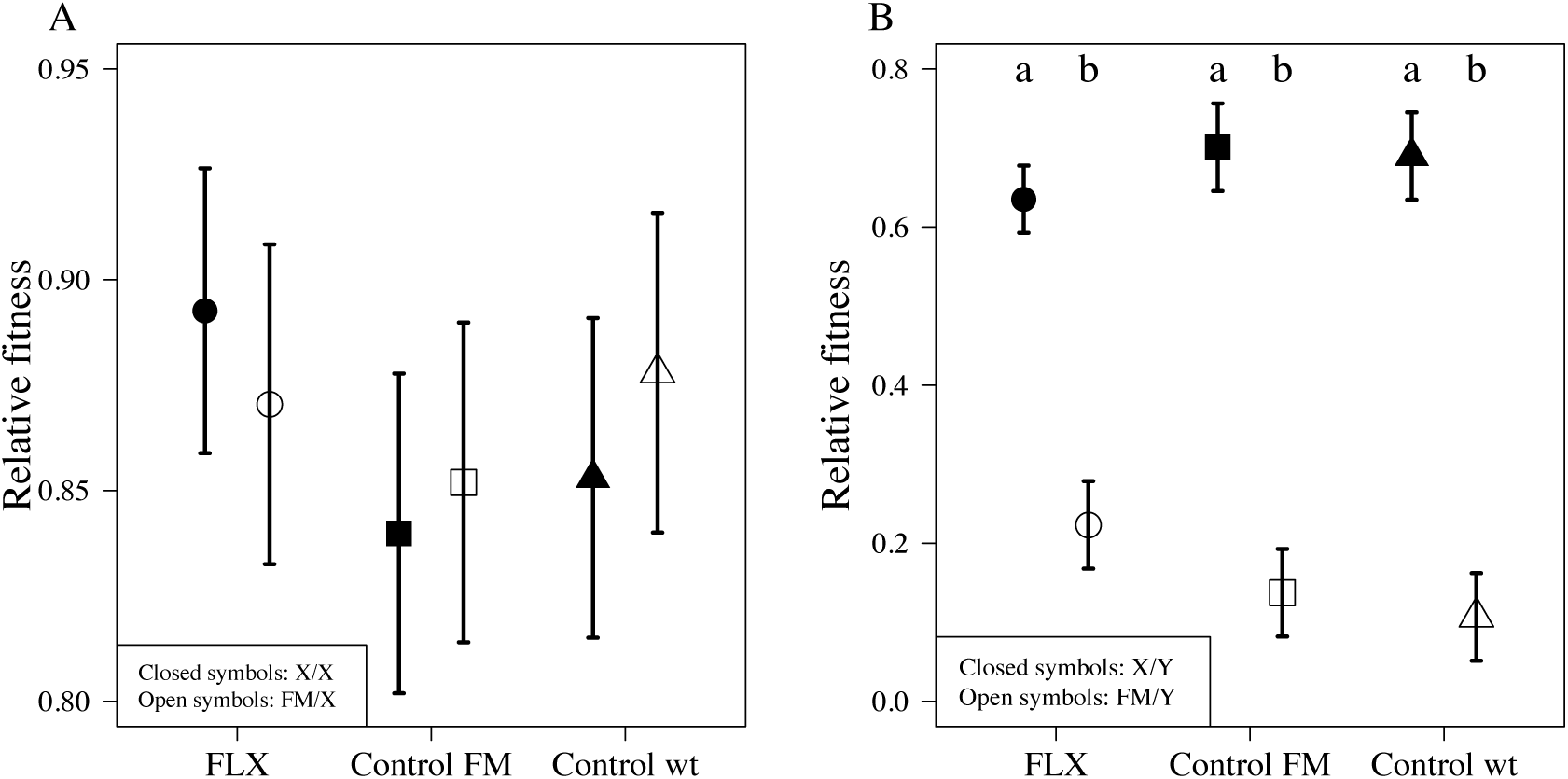
Relative fitness for: (A) females at generation 127 and (B) males at generation 127. Closed symbols: X/X or X/Y, open symbols: FM/X or FM/Y. FLX: circle, control FM (CFM): square, and control wild-type (Cwt): triangle. Mean (±SE) average across the four replicate populations. Letters indicates significance (Tukey HSD; *P* < 0.05).

## Discussion

### General patterns and caveats

We hypothesised that enforcing female-limited evolution of the X chromosome would result in a feminized phenotype in traits assumed to be sexually antagonistic. Consistent with our expectations, we found that FLX flies increased in size relative to Cwt, while also developing faster, both an indication of a more feminized phenotype in both sexes. Contrary to our expectations, there was no evidence of sexual antagonism for fitness, and results for locomotory activity were most consistent with an effect of the FM balancer.

Long and Rice (2007) found that adult locomotory activity is the target of intralocus sexual conflict, with a positive correlation between male fitness and activity level but a negative correlation for females. As less active females are courted less frequently by males (Tompkins et al. 1982), it follows that females would benefit from reduced locomotory activity, when male courtship is harmful to females (Nandy et al. 2013). We found that females overall were significantly less active than males, which is consistent with the assumption that it is beneficial for females to be less active. Abbott *et al*. (2019) showed that after ∼30 generations of an MLX evolution experiment, the locomotory activity of MLX females had increased to levels comparable to that of males. We did find that FLX females were less active than FLX males, but there was no evidence that FLX males had decreased locomotion activity compared to the other treatments. In contrast, the CFM males seemed to have the lowest activity. We therefore speculate that the lack of impact on male locomotion activity in FLX could be due to a cancelation of effects between selection and the adaption to the FM balancer (discussed in more detail below).

Unlike previous ML experiments (Prasad et al. 2007; Abbott et al. 2010) we were not able to find a significant effect of selection on reproductive fitness in the FLX flies. There are several plausible explanations for the lack of response in fitness to the selection regime. First, we used a different experimental setup, as we selected only on the X chromosome while previous authors selected on all major chromosomes. Though theory predicts that the X chromosome should be enriched for sexually antagonistic variance (Rice 1984), it is not limited to the X (Fry, 2010) and a recent genome-wide association study did not support the idea of the X as a hot spot for sexually antagonistic variance in this species (Ruzicka et al. 2019). So, limiting selection to one sex at all genomic loci simultaneously, as in the ML experiments, may have provided more power to detect sexually antagonistic effects. Abbott *et al*. (2019) carried out a male-limited X chromosome evolution experiment, and were also not able to find an equal response to selection as previous ML evolution experiments. Though they did find an increase in male reproductive fitness, they did not find an equivalent decrease in female fitness. They discussed that the lack of decrease in female fitness might be due to dominance effects on the X chromosome, and this explanation is plausible in this case as well. Because the X chromosome is hemizygous in males, any male-beneficial recessive loci will be exposed to selection in males, even if detrimental to females, but will be hidden from selection in females because they have two X chromosomes. On the other hand, any female-beneficial dominant loci will be expose to selection in females, i.e. the dominance effect (Rice 1984; Patten 2018). So if female-benefit X-linked loci related to fecundity are mostly dominant on the X, then females may already be close to their optimum for this trait. We are currently preparing a more in-depth analysis of possible dominance and epistatic effects in these populations to investigate this explanation.

A second non-exclusive explanation for the lack of response in fitness is weak selection in females compared to males. Male traits are predicted to be more evolvable than female traits (Connallon and Knowles 2005; Allen et al. 2018), and the X chromosome is expected to experience stronger selection in males, as they are hemizygous (Patten 2018). Therefore, the lower response we found in the evolution experiment compared to the ML experiments could be due to males experiencing stronger selection on the X chromosome than females, and that male traits are more likely to change as a result of experimentally enforced sex-limited selection over any given evolutionary timespan.

Finally, adaptation to the FM balancer chromosome might also have obscured phenotypic responses to selection in the FLX lines. An analysis of gene expression differences between treatments suggests that this is the case, as there is evidence of some differentiation between the FLX and CFM expression profiles (Lund-Hansen 2017).

### Effect of the balancer chromosome

The CFM treatment was devised to control for any unknown effects of adaptation to the FM balancer. If the FM balancer functioned like a normal X chromosome, phenotypic traits of CFM flies should be similar to the Cwt treatment. However, it was clear from the phenotypic assays that the CFM flies were not always similar to the Cwt flies, and that carrying the FM balancer did have an effect on the flies. The long-term effect of carrying the FM balancer was particularly evident when comparing the results for body size and reproductive fitness in the early generations to the later ones. Reproductive fitness was greater in CFM males in the later generations, as well as CFM females increasing in body size to achieve the same size as the FLX females. These results indicate that there was long-term adaptation to the FM balancer in CFM flies, which makes it difficult to separate the effect of the FLX selection from adaptation to the FM balancer in the FLX treatment as well. We speculate that the inconsistency between the early and late phenotypic results could be due to selection for male-beneficial genes on the autosomes to ameliorate the negative effect of carrying the FM balancer in males. So when FLX males express the FLX-selected X chromosome this could lead to conflict between selection for female-beneficial alleles on the X chromosome and male-beneficial alleles on the autosomes.

Another side-effect of using the FM balancer is that it lowers the quality of the FM/Y males. Lower quality males should relax sexual selection (Holland and Rice 1999; Pitnick et al. 2001; Morrow et al. 2008; Nandy et al. 2013; Innocenti et al. 2014), so in addition to the removal of intralocus conflict, the presence of the balancer might also have reduced interlocus conflict. This could explain why for some traits CFM flies changed in the same way as was predicted for FLX flies under release from counter-selection in males (e.g. development time, female fitness at generation 39/41).

We are currently testing these hypotheses about the effect of the balancer using gene expression data (Lund-Hansen 2017) and carrying out a quantitative genetic analysis of the selection lines, to disentangle effects resulting from different selection pressures on the X versus the autosomes.

### Trade-offs between traits

An easily observed phenotypic trait in *D. melanogaster* is body size, which has been positively correlated to female fecundity (Robertson 1957). Abbott *et al*. (2010) showed that limiting selection to males caused body size to decrease in both sexes, which would be harmful to female fecundity. We therefore expected that limiting selection to females would result in increased body size in both sexes, which was true at both generations 18 and 72. However, unlike the expected detrimental effect to female fecundity of a smaller body size, there are conflicting results as to where the optimal male body size lies. Evolution experiments indicate that the male optimum for body size is to be small (Pitnick et al. 2001; Prasad et al. 2007; Abbott et al. 2010; Pischedda et al. 2012) whereas studies on natural populations show that being large is advantageous (Partridge et al. 1987a,b). These conflicting results suggest that selection on male body size is context-dependent, and may not be subject to intralocus sexual conflict to the same extent in the wild.

As females develops faster than males (Bonnier 1926), it suggest that there could be selection on females for faster development time. We did indeed find that FLX flies had a decreased development time compared to Cwt flies. It can seem counterintuitive that the larger FLX flies also develop faster, as faster development time normally is found to decrease size (Prasad et al. 2000). However, early hatching has been positively correlated with a larger egg output for females as well as size (Robertson 1957), so for female fitness, developing fast and large would be beneficial. However, it is not likely to be beneficial for males to develop fast and large, as previous results suggest there is a trade-off between body size and gonad development (Bonnier 1926; Nunney 1996). This trade-off could explain why we did not find a correlated increase in male reproductive fitness associated with the larger body size in FLX males, as they could be experiencing a trade-off between faster development time and investment in gonad development. This trade-off is likely to be exacerbated by the effect of the FM balancer in reducing the intensity of sexual conflict. In line with this idea, Prasad *et al*. (2007) found that smaller males had an increase in development time and reproductive fitness, which could indicate that ML males had more energy (and time) to invest in gonad development. There is also some evidence that FLX males perform better in mating success but more poorly in sperm competition (Manat et al. unpublished data), suggesting a trade-off between these traits. Collectively these results indicate that there is an on-going sexual conflict over development time in *D. melanogaster*.

## Conclusion

Previous evolution experiments have shown that it is possible to eliminate intralocus sexual conflict by limiting selection to one sex, and thereby removing counter selection. These previous evolution experiments showed responses that shifted numerous phenotypic traits towards the male optimum, causing harm to females that expressed the male-limited genome. However, these previous studies have been exclusively male-limited, restricting the generality of these tests. To determine if the response to limited selection would be the same in a female-limited experiment, we performed the first female-limited X chromosome evolution experiment. We did not find a strong response to selection in reproductive fitness or locomotory activity, but we did find a response in body size and development time. These results suggest that body size and development time are likely experiencing ongoing sexually antagonistic selection, and that a substantial proportion of the genetic variance for these traits must be X-linked. Also, adaptation to the FM balancer appears to have had an effect on the flies, so further investigation is necessary to determine how compensatory alleles on the autosomes and reduced interlocus sexual conflict may contribute to the changes seen here. In conclusion, as the first female-limited X chromosome evolution experiment, this study adds to the growing list of evidence for sexually antagonistic variance in the genome and how it may constrain the evolution of sexual dimorphism.

## Acknowledgments

Qinyang Li, Claire Webster, and Ilona Flis for technical help throughout the experiment. A number of bachelor students at University of Sussex and Lund University for help with data collecting. This work was supported by a Swedish Research Council grant to KKL-H, ERC grants to JKA and EHM, and a Royal Society University Research Fellowship to EHM. The authors have no conflict of interest to declare.

