## Supplementary material for "Feminisation of complex traits in *Drosophila melanogaster* via female-limited X chromosome evolution"

Table S1: Summary of the t-tests between the two FLX female types and between the two control FM (CFM) female types.

| Source | *df* | t | *P* |
|---|---|---|---|
| Fitness, generation 15 & 18 | | | |
| FLX 1X vs. 2X | 75.33 | 1.53 | 0.13 |
| CFM 1X vs. 2X | 75.98 | -0.51 | 0.61 |
| Fitness, generation 39-41 | | | |
| FLX 1X vs. 2X | 85.19 | 1.22 | 0.23 |
| CFM 1X vs. 2X | 90.28 | 2.21 | *0.03* |
| Thorax size, generation 15 & 18 | | | |
| FLX 1X vs. 2X | 239 | 2.96 | $3.39e^{-03}$ |
| CFM 1X vs. 2X | 254.96 | -0.38 | 0.71 |
| Thorax size, generation 72 | | | |
| FLX 1X vs. 2X | 156.92 | -3.54 | $5.26e^{-04}$ |
| CFM 1X vs. 2X | 146.04 | $2.16e^{-08}$ | 1 |
| Development time, generation 43 | | | |
| FLX 1X vs. 2X | 35.72 | 0.44 | 0.66 |
| CFM 1X vs. 2X | 37.99 | -0.05 | 0.96 |
| Locomotion, generation 123 | | | |
| FLX 1X vs. 2X | 78 | 0 | 1 |
| CFM 1X vs. 2X | 77.30 | 0 | 1 |

Table S2: Summary of the results from ANOVA analysis of linear mixed models

| Source | *df* | F | $\chi^2$ | *P* |
|---|---|---|---|---|
| Fitness, generation 15 & 18 | | | | |
| Treatments | 2 | 0.21 | | 0.81 |
| Sex | 1 | 0.05 | | 0.82 |
| Treatments × Sex | 2 | 1.74 | | 0.18 |
| Fitness, generation 39-41 | | | | |
| Treatments/Types | 7 | 1.39 | | 0.48 |
| Thorax size, generation 15 & 18 | | | | |
| Treatments/Types | 7 | 361.64 | | $< 2.2e^{-16}$ |
| Thorax size, generation 72 | | | | |
| Treatments/Types | 7 | 79.94 | | $4.43e^{-15}$ |
| Development time, generation 43 | | | | |
| Treatments | 2 | 3.70 | | 0.06 |
| Sex | 1 | 3.37 | | 0.07 |
| Treatments × Sex | 2 | $8.90e^{-03}$ | | 0.99 |
| Locomotion, generation 123 | | | | |
| Treatments | 2 | | 21.26 | 0.53 |
| Sex | 1 | | 9.92 | $1.64e^{-03}$ |
| Treatments × Sex | 2 | | 3.22 | 0.20 |
| FM female fitness assay, generation 127 | | | | |
| Treatments | 5 | 0.65 | | 0.66 |
| FM male fitness assay, generation 127 | | | | |
| Treatments | 5 | 39.93 | | $3.98e^{-09}$ |
